# From nest to hatchling: nidobiome assembly and host selection in oviparous vertebrates without parental care

**DOI:** 10.64898/2026.05.26.727962

**Authors:** Ana Sofia Carranco, David Romo, Maria de Lourdes Torres, Kerstin Wilhelm, Simone Sommer

**Affiliations:** Institute of Evolutionary Ecology and Conservation Genomics, University of Ulm, Ulm, Germany; Tiputini Biodiversity Station, Universidad San Francisco de Quito, Cumbaya–Quito, Diego de Robles y Via Interoceanica s/n, Quito, 170901, Ecuador; Laboratorio de Biotecnología Vegetal, Universidad San Francisco de Quito USFQ, Diego de Robles y Via Interoceanica s/n, Quito, 170901, Ecuador

**Keywords:** early microbial assembly, host-filtering, priority effects, nidobiome, oviparous vertebrates, freshwater turtles

## Abstract

Early microbial assembly is fundamental for host development and fitness, yet in oviparous vertebrates lacking parental care, the mechanisms by which eggs and neonates acquire their microbiome remain poorly understood. The nidobiome concept predicts that maternal and environmental sources, and host selection, collectively shape early microbial communities. To test this, we conducted a longitudinal study on microbial assembly in the yellow-spotted Amazon river turtle (*Podocnemis unifilis*). We sampled maternal mucosa, freshly laid outer eggshells, and nest sand before incubation, along with hatchery sand, outer and inner eggshells, and hatchling cloaca after incubation. Using 16S and ITS amplicon sequencing to quantify microbial diversity, we observed that freshly laid outer eggshells shared bacterial diversity with maternal mucosa, whereas nest sand harboured a distinct community, providing evidence of early vertical transmission. After incubation, eggshell microbial communities became more even without a change in richness, while nest sand communities exhibited the opposite pattern, reflecting divergent host-environment assembly dynamics. SourceTracker analyses confirmed contributions from both maternal and environmental sources, yet inner eggshells and hatchling cloaca assembled distinct communities compared with outer eggshells, indicating that host selection is largely deterministic. Interkingdom interactions favoured beneficial microorganisms, suggesting active host filtering throughout development. Even in the presence of fusariosis, a widespread threat in turtle nests, host selection enriched protective taxa, linking microbiome composition to hatching success. These findings demonstrate that maternal seeding, priority effects, and deterministic host selection govern early-life microbiome assembly in oviparous reptiles without parental care, with direct implications for microbe-informed conservation strategies.

**Importance:** Understanding early-life microbial assembly is critical for uncovering the ecological and evolutionary drivers of host-microbiome interactions. This study reveals how microbial communities develop in oviparous vertebrates lacking parental care, highlighting key mechanisms such as vertical transmission, host selection, and environmental modulation that shape early-life microbiomes. Our results demonstrate that maternal microbiota seeding and priority effects play fundamental roles in structuring egg-associated microbial communities, directly influencing host fitness and survival. By identifying deterministic processes that enrich beneficial microorganisms during early development, this research underscores the pivotal role of microbiomes in host resilience and adaptation. These insights carry urgent conservation implications, offering microbe-informed strategies to mitigate emerging threats, such as diseases in vulnerable species like the yellow-spotted Amazon river turtle, and emphasising the broader importance of microbiome stewardship for ecosystem health.

## Introduction

The microbiome has emerged as a fundamental component of animal biology, with profound influences on host development, immunity, disease resistance, and overall health (1–4). A critical step in establishing host-microbiome relationships is the initial acquisition of microbial communities. In many animals, including humans and oviparous vertebrates with parental care, microbial acquisition occurs through vertical (maternal) and horizontal (environmental) transmission, and extended parental contact enhances continuous microbial colonisation (5–10). In contrast, in oviparous species lacking parental care, maternal microbial transfer is restricted to internal egg development and oviposition, and post-hatching microbiome assembly is therefore likely dominated by environmental sources, as demonstrated in turtles (11). Nevertheless, studies often consider vertical and horizontal transmission as independent processes, overlooking their combined effects and the functional significance of their relative contributions to microbiome assembly.

The nidobiome concept offers an integrated framework that recognises microbiome assembly as a dynamic process, in which maternal and environmental microbial inputs, combined with host selection and priority effects, jointly shape offspring microbiome development and fitness (12, 13). Although this framework has typically been applied to species with parental care, where mothers actively modify the nidobiome and protect eggs from bacterial and fungal infections through behaviours such as nest sanitation or antimicrobial secretions (14), species lacking post-oviposition maternal influence depend on initial microbial provisioning during oviposition and nest-site selection (15). Recent work on lizards suggests that vertically transmitted bacterial communities can protect eggs against fungal and bacterial infections (16), while studies from freshwater turtles indicate that bacterial communities are established through mixed-mode transmission (17). Nevertheless, it remains unclear how environmental challenges, particularly pathogen exposure, interact with vertical and horizontal microbial inputs to shape microbiome assembly and influence offspring fitness in species without parental care.

Environmental stressors, especially pathogenic exposure, can profoundly alter host-associated microbiomes, with outcomes ranging from enhanced resistance to dysbiosis (18–20). For oviparous vertebrates lacking parental care, this challenge is particularly critical during incubation, when all microbial assembly occurs within the egg and nest environment, and offspring cannot rely on parental intervention to buffer against pathogens. Disruptions to microbial dynamics at this crucial stage can have cascading effects on disease resistance, development, and overall population health (21). Yet, the mechanisms governing nidobiome resilience remain poorly understood, including which microbial lineages confer protection, how priority effects and host selection influence pathogen susceptibility during incubation, and how maternal microbial provisioning shapes offspring disease resistance in the absence of post-oviposition care.

The yellow-spotted Amazon river turtle (*Podocnemis unifilis*) provides an ideal system to investigate these knowledge gaps. This species lacks parental care and is vulnerable to egg fusariosis, a fungal infection caused by the *Fusarium solani* species complex (FSSC) that results in turtle hatchling mortality and population declines (22–24). Previous work has shown that *P. unifilis* eggs harbour an inner microbiome, particularly a fungal community, that may provide antifungal protection and enhance hatching success (11, 25). However, the mechanisms governing the acquisition and assembly of disease-resistant bacteriomes and mycobiomes remain unclear. Specifically, it is unknown which microbial lineages are maternally provisioned versus environmentally acquired, how priority effects and host selection shape microbial community assembly during incubation, and whether fusariosis disrupts these processes. Using the nidobiome framework, in the present study, we investigated the mechanisms underlying early-life microbiome assembly and its role in disease resistance.

We hypothesised that the nidobiome plays a crucial role in promoting disease resistance in species lacking parental care, involving four interconnected processes during incubation. First, priority effects influence microbial community composition: early-colonising microbes, acquired both vertically and horizontally, establish dominance and persist in abundance even when alternative taxa are present. Second, host selection shapes microbial communities through deterministic processes, whereby the host selectively retains beneficial microorganisms, leading to a microbiome that converges toward a functional, disease-resistant state and shows reduced dispersion compared to the surrounding environment. Third, specific bacterial and fungal taxa from maternal and environmental sources are retained by the host, although this pattern can be disrupted by fusariosis infection. Finally, interkingdom interactions contribute to microbiome resilience: bacterial-fungal relationships formed during incubation persist after hatching, providing antifungal protection even in the presence of *Fusarium* pathogens. Collectively, these processes highlight how incubation-stage microbiome assembly, guided by the nidobiome framework, can protect offspring from fungal pathogens during a vulnerable developmental window in the absence of parental care.

## Results

### Sample overview and FSSC infection patterns

We successfully obtained bacteriome sequences from 490 samples and mycobiome sequences from 239 samples (Table S1). FSSC infection was detected exclusively in post-incubation samples, suggesting that pathogen colonisation occurs during incubation. Of the 11 investigated nests, 10 contained FSSC-infected samples. Infection prevalence varied by sample type: 54% of hatchery sand samples (n = 6/11), 61% of outer eggshell samples (n = 46/75), 50% of inner eggshell samples (n = 38/76), and 15% of hatchling cloaca samples (n = 10/65) were positive for FSSC pathogens (Table S1). FSSC identity was confirmed by sequencing TEF-1α amplicons from representative positive samples, with *Fusarium keratoplasticum* being the dominant pathogen.

### Changes in bacteriome and mycobiome alpha diversity pre- and post- incubation, and effects of Fusarium infections

The nidobiome concept posits that interactions among parents, offspring, and the nest environment play a key role in the initial assembly of host microbial communities. To assess whether this concept applies to oviparous vertebrates lacking parental care and whether this process is influenced by pathogens, we analysed alpha diversity in samples collected before (pre-incubation) and after (post-incubation) incubation. These samples represented nidobiome source components (maternal mucosa, nest sand) and the developmental trajectory from maternal and environmental provisioning to offspring microbiota establishment (outer eggshells, inner eggshells, hatchling cloaca). Linear models were used to evaluate shifts in microbial diversity and the effects of fusariosis infections on community communities.

We observed significant variation in bacteriome and mycobiome species richness and Shannon diversity across sample types throughout the incubation period (Table S2). During the pre-incubation phase, there were no significant differences in bacteriome species richness between maternal mucosa and outer eggshells (Fig. 1a), but a small significant difference was observed in bacteriome and mycobiome Shannon diversity and mycobiome richness (p < 0.001, Fig. 1b, 1c, 1d; Fig. S1). Conversely, nest sand exhibited a greater bacteriome diversity compared to both maternal mucosa and outer eggshells (p < 0.001, Fig. 1a, 1b, Fig. S1). However, no significant differences were observed in mycobiome diversity and richness between the two components of the nidobiome: maternal mucosa and the nest environment (Fig.s 1c, 1d; Fig. S1). These findings suggest that the maternal inoculum is transmitted vertically from the maternal cloacal mucosa to the outer eggshell and that the microbial availability in the nest environment further contributes to the egg microbiome prior to incubation.

**Fig. 1.**
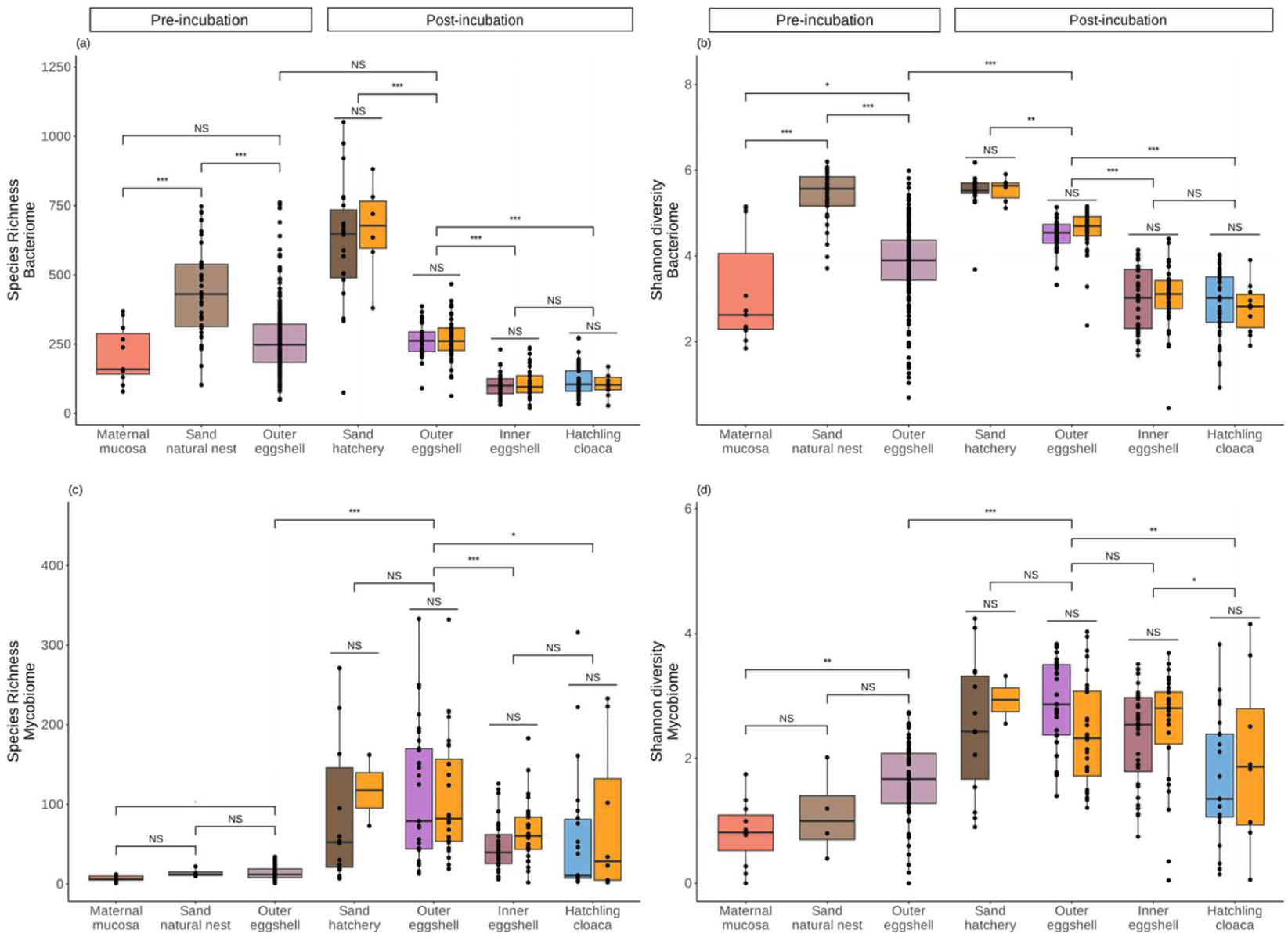
Changes in bacteriome and mycobiome alpha diversity during incubation and effects of *Fusarium* infections. Species richness and Shannon diversity index of the bacteriome (a, b) and mycobiome (c, d) are shown in boxplots comparing pre-incubation environmental (sand from the natural nest) and biological samples (maternal mucosa, outer eggshell), and post-incubation environmental (sand hatchery) and biological samples (outer eggshell, inner eggshell, hatchling cloaca), alongside fusariosis infection status. Tukey’s post hoc test with multiple comparisons corrections for general linear models was used to assess differences between pre- and post-incubation samples and the effect of fusariosis infection. Significant differences are shown with an (*) according to the adjusted p-value (“***” <0.001; “**” <0.01; “*” <0.05; “.” <0.1), and groups with no significant differences (>0.1) are indicated by the acronym N.S. (not significant).

Species richness and Shannon diversity of both the bacteriome and mycobiome in the nest environment (sand) and on outer eggshells increased significantly from pre- to post-incubation (p < 0.001, Fig. 1, Table S2). Outer eggshells, which are continuously exposed to nest microbes during incubation, showed parallel increases in bacteriome and mycobiome Shannon diversity and in mycobiome richness post-incubation (p < 0.001, Fig. 1), whereas bacteriome richness remained stable (Fig. 1a). These findings suggest that during incubation, the maternal inoculum is effectively assembled on the outer eggshell, with the nest environment playing an active role in microbial colonisation and eggshell microbial assembly. In contrast, post-incubation inner eggshells and hatchling cloaca showed markedly reduced diversity. Both bacteriome and mycobiome species richness and bacteriome Shannon diversity were significantly lower in the inner eggshell and hatchling cloaca than in the outer eggshell (p < 0.001, Fig. 1a- c). Additionally, mycobiome Shannon diversity of the hatchling cloaca was lower than in both outer and inner eggshell samples (p < 0.001, Fig. 1d, Fig. S1), suggesting active host-mediated microbial filtering.

FSSC infection had no significant effect on the nest or eggshells and hatchling’s cloaca microbiome alpha diversity, as neither infection status nor its interaction with sample type influenced bacteriome or mycobiome alpha diversity (Fig. 1; Tables S3 and S4). This suggests that nidobiome assembly processes and the hosts’ selection of microbial communities remain robust even in the presence of fungal pathogens.

### Microbial community composition during pre- and post- incubation and effects of *Fusarium* infections

To examine shifts in community composition across the nidobiome-offspring continuum, we performed redundancy analyses (RDAs) using centred log-ratios (rCLR) to account for the compositional nature of microbiome data. Sample type explained significant variation in both bacterial (F_8,490_ = 8.975; p < 0.001; adjR² = 11%, Fig. 2a, Table S5) and fungal composition (F_6,232_ = 2.508; p < 0.001; adjR² = 3%, Fig. 2f, Table S6) across pre- and post-incubation samples, with strong differences observed between all sample types before and after incubation (Table S7a, Table S8a). These compositional shifts demonstrate that nidobiome microbial communities are temporally dynamic and that distinct microbial assemblages characterise different nidobiome components (nest sand, maternal mucosa) and host microbial assembly.

**Fig. 2.**
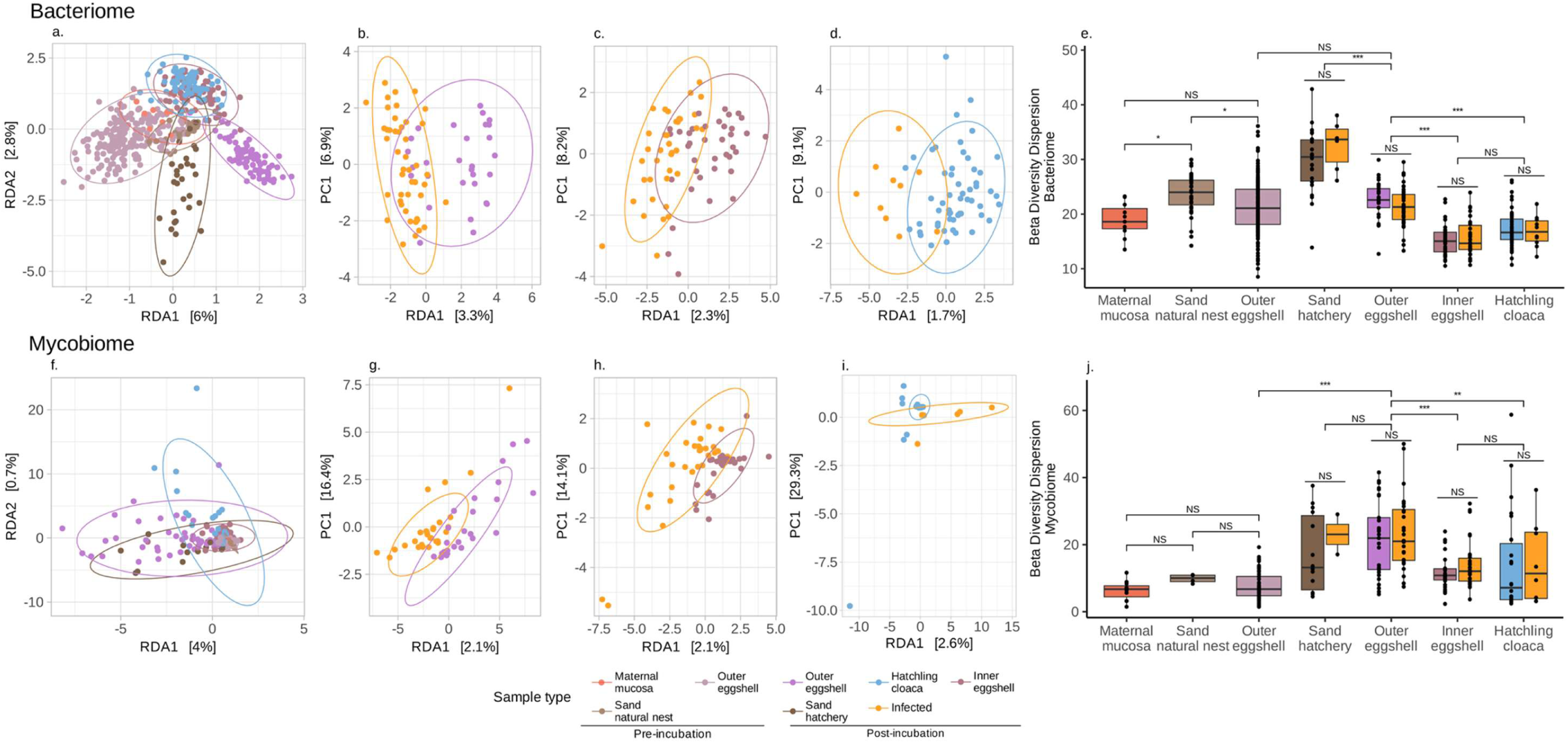
Changes in bacteriome and mycobiome composition during pre- and post- incubation and effects of fusarium infections. Redundancy analysis (RDA) ordination plots, using Euclidean distances, show the beta diversity of bacteriomes (upper panels) and mycobiomes (lower panels) according to: (a, f) sample type pre-incubation (sand from the natural nest, maternal mucosa, and outer eggshell) and post-incubation (sand hatchery, outer eggshell, inner eggshell, and hatchling cloaca); (b, c, d, g, h, i) FSSC infection status in post-incubation samples (infected: orange); (e, j) box plots of community dispersion with Tukey’s multiple comparison test for GLS models. Significant differences are shown with an (*) according to the adjusted p-value (“***” <0.001; “**” <0.01; “*” <0.05; “.” <0.1), and groups with no significant differences (>0.1) are indicated by the acronym N.S. (not significant).

Microbial composition mirrored the pattern of reduced diversity observed in inner eggshells in the alpha diversity analyses. RDA analyses showed particularly strong differences between outer and inner eggshells, as well as between inner eggshell and hatchling cloaca samples (Table S7a, Table S8a), suggesting that the host selects for specific microbial taxa from the nidobiome as microbes move into the inner eggshell.

Fusariosis infection explained only a small proportion of the variation in bacteriome community composition (F_1,242_ = 2.709; p < 0.001; adjR² = 0.7%, Fig. 2b, c, d, Table S5b, c), with effects being most pronounced in the outer and inner-eggshell samples (Table S7c). Notably, no effect of FSSC infection was observed in the hatchling cloaca, suggesting that host filtering mechanisms that selectively retain beneficial nidobiome microbes may simultaneously exclude or limit pathogen colonisation in offspring tissues. Furthermore, we observed no effect of FSSC infection on mycobiome composition in biological samples (F_1,163_ = 1.134; p = 0.196; adjR² = 0.08%, Fig. 2g, h, i, Table S6b) or in sand environment samples. Consistent with our previous reports on inner eggs (25), our results extend this pattern to the hatchling cloaca microbiome, showing that pathogen presence does not disrupt host selection processes shaping offspring microbial communities.

### Homogeneous inner-egg and hatchling microbiomes reflect host filtering rather than infection

To determine whether offspring microbiome assembly from the nidobiome follows deterministic (host-driven selection toward consistent communities) or stochastic (random acquisition resulting in variable communities) processes, we examined beta diversity dispersion (community heterogeneity) across sample types using generalised least squares (GLS) models.

Community dispersion of both the bacteriome and mycobiome was significantly explained by sample type (bacteriome: F_6,482_= 66.682; p < 0.001, adjR²= 44%, Fig. 2e; mycobiome: F_6,231_= 15.557; p < 0.001, adjR²= 29%, Fig. 2j; model selection Table S9), but not by infection status (bacteriome: F_1,482_= 0.005; p = 0.942; mycobiome: F_1,231_= 1.653; p = 0.200). Nidobiome environmental samples (sand from natural nests and hatcheries) exhibited significantly greater heterogeneity than all biological samples (p < 0.001, Fig. S2). Heterogeneity further increased after incubation in the nest environment (p < 0.001, Fig. S2), suggesting that the microbial communities in the nest change during incubation, resulting in a diverse pool of microbial available to the clutch.

In contrast, bacteriome and mycobiome heterogeneity progressively decreased from the outer eggshell to the inner eggshell and hatchling cloaca (p < 0.001, Fig.2), with no significant difference observed between the inner eggshell and hatchling cloaca, suggesting that host selection during embryonic development establishes a homogeneous microbial community that is maintained in the hatchlings’ cloaca. Overall, these findings support a deterministic role for host selection over stochastic assembly, whereby hosts selectively filter specific microbes from heterogeneous nidobiome sources (variable nest environments and maternal provisioning), resulting in relatively homogeneous offspring microbiomes. The inner-egg bacterial community appears to form the foundation of the hatchling cloacal bacteriome, as previously demonstrated in this turtle species (11).

### Contribution of maternal and environmental sources in shaping egg and hatchling microbiomes with infection-dependent variation

To determine whether maternal and environmental sources contribute to egg and hatchling microbiome assembly, and whether fusariosis infection disrupts this process, we used SourceTracker2 to trace bacterial and fungal genera from source (maternal cloaca and nesting sand) to the “sink” outer- and inner-eggshells and hatchling cloaca.

Both maternal and environmental sources contributed bacterial and fungal symbionts to the hatchling cloaca. Uninfected samples shared 41 bacterial (Fig. 3a) and 15 fungal genera (Fig. 3c) between the two sources in the hatchling cloaca. In contrast, FSSC-infected samples shared 36 bacterial (Fig. 3b) and five fungal genera (Fig. 3d) from both sources in the hatchling cloaca. These results suggest that both maternal and environmental sources contribute to the microbiome of the egg and hatchling cloaca.

**Fig. 3.**
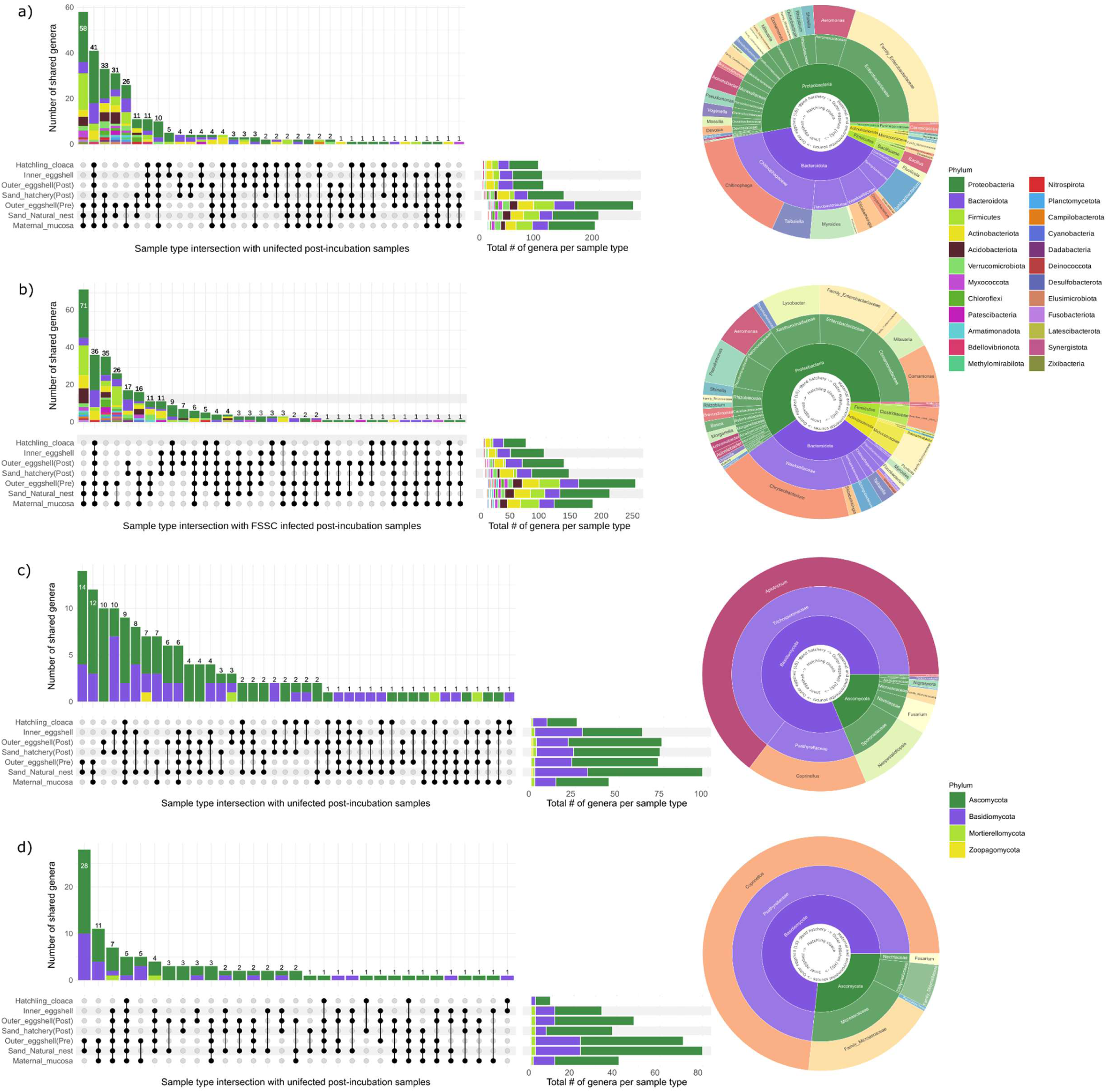
Microbial transmission patterns from maternal and environmental sources across developmental stages in FSSC-uninfected and infected samples. Results are shown for the bacteriome in (a) uninfected, (b) infected samples, and for the mycobiome in (c) uninfected, (d) infected samples. Left-hand panels display UpSet plots illustrating intersections of genera across pre-incubation (maternal mucosa, sand natural nest, outer-eggshell) and post-incubation sample types (sand hatchery, outer-eggshell, inner-eggshell, hatchling cloaca). Horizontal bars indicate the number of genera detected per sample type, while vertical bars represent intersection sizes (genera shared among specific sample combinations). Right-hand panels show sunburst plots depicting transmission pathways of core genera (present across all sample types) from combined source environments (maternal mucosa and sand natural nest) through successive developmental stages. The inner ring represents sample pathways, and outer rings indicate hierarchical taxonomic classification (phylum → family → genus). Colours denote phylum-level (middle rings) and genus-level (outermost rings) taxonomy, and are consistent across all panels.

Several bacterial genera sourced from the maternal cloaca and the nest environment showed reduced abundance in FSSC-infected samples, including *Taiabella*, *Stenotrophomonas*, *Sphingobacterium*, *Pseudomonas*, *Flavobacterium*, *Family Enterobacteriaceae*, *Elizabethkingia*, and *Chitinophaga* (Fig. 4a). Three genera, *Paenathrobacter*, *Morganella*, and *Bosea,* were only shared across samples when infection was present (Fig. 4a). Additionally, *Comamonas*, *Chryseobacterium*, *Clostridium sensu stricto 1,* and *Lysobacter* showed increased abundance in FSSC-infected samples (Fig. 4a). Among the fungal genera sourced from the maternal cloaca and the nest environment, *Apiotrichum* and *Neopestalotiopsis* were exclusively present in uninfected samples (Fig. 4b). In contrast, Family Didymellaceae was only detected in FSSC-infected samples (Fig. 4b). Notably, both *Coprinellus* and *Fusarium* showed higher abundance in FSSC-infected samples compared to uninfected samples, with *Coprinellus* appearing to originate from the sand environment, while *Fusarium* originated from both the maternal and environmental sources and showed increased abundance after incubation from the maternal source (Fig. 4b). Our results provide strong evidence for maternal and environmental contributions to the eggshell and cloacal microbiomes during incubation. Additionally, the impact of FSSC infection on the abundance of several taxa suggests that microbial transmission dynamics, in conjunction with host selection, may play a crucial role in infection resistance and successful hatching.

**Fig. 4.**
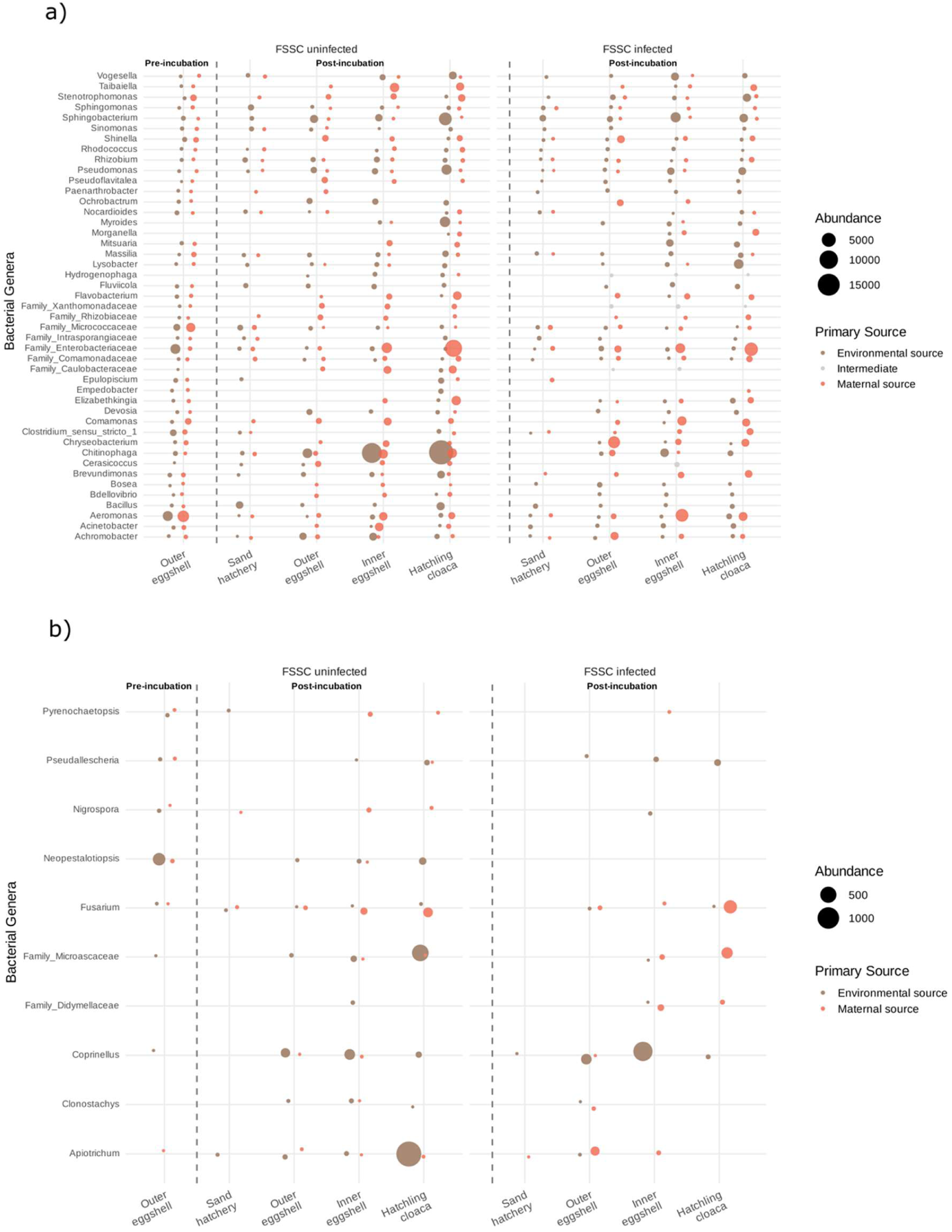
Bubble plots illustrating source-specific bacteria and fungal genera shared from the two primary sources across all the sample types in FSSC-uninfected and infected samples. The distribution of key bacterial (a) and fungal (b) genera identified from intersection analyses is shown. Bubble sizes indicate the relative frequency of each genus within a given sample type, while bubble colour denotes the maternal (maternal mucosa represented with peach colour) and environmental source (sand natural nest represented with brown colour). Sample types are arranged chronologically from left to right across the pre- and post-incubation periods. Only genera from major UpSet intersections are displayed, representing the core shared microbiome across development.

### Priority effects and host filtering enrich functionally relevant microbes during incubation

During incubation, the microbiome assemblage is influenced by microbial priority effects, in which initial microbial communities occupy niches and prevent the establishment of other taxa. We tested whether priority effects are at play for nidobiome microbial sources by performing an analysis of composition of microbiomes with bias correction (ANCOM-BC2). We compared nidobiome vertical (maternal mucosa) and horizontal sources (sand natural nest) with pre- (outer eggshell) and post-incubation samples (sand hatchery, outer eggshell, inner eggshell and hatchling cloaca), and tested for the effect of FSSC-infection in post-incubation samples.

We identified 46 bacterial genera (q-value < 0.005, Fig. 5a) to be highly abundant in the different samples and six fungal genera enriched in the post-incubation samples but not in the hatchling cloaca (q-value < 0.05, Fig. 5d). These included the bacterial genus *Sphingobacterium*, *Devosia*, *Taiabella*, *Shinella*, *Chitinophaga*, *Bosea*, *Bdellovibrio*, *Hydrogenophaga*, Family Xanthomonadaceae, *Pseudoflaviatella*, *Brevundimonas*, Family Rhizobiaceae, *Sphingomonas*, *Lysobacter*, Family Comamonadaceae, *Cerasicoccus*, *Achromobacter*, *Fluviicola*, *Flavobacterium* and Family Caulobacteraceae; and the fungal genus *Apiotrichum*, *Coprinellus*, *Pyxiodophora*, *Cladosporium* and Family Didymellaceae.

**Fig. 5.**
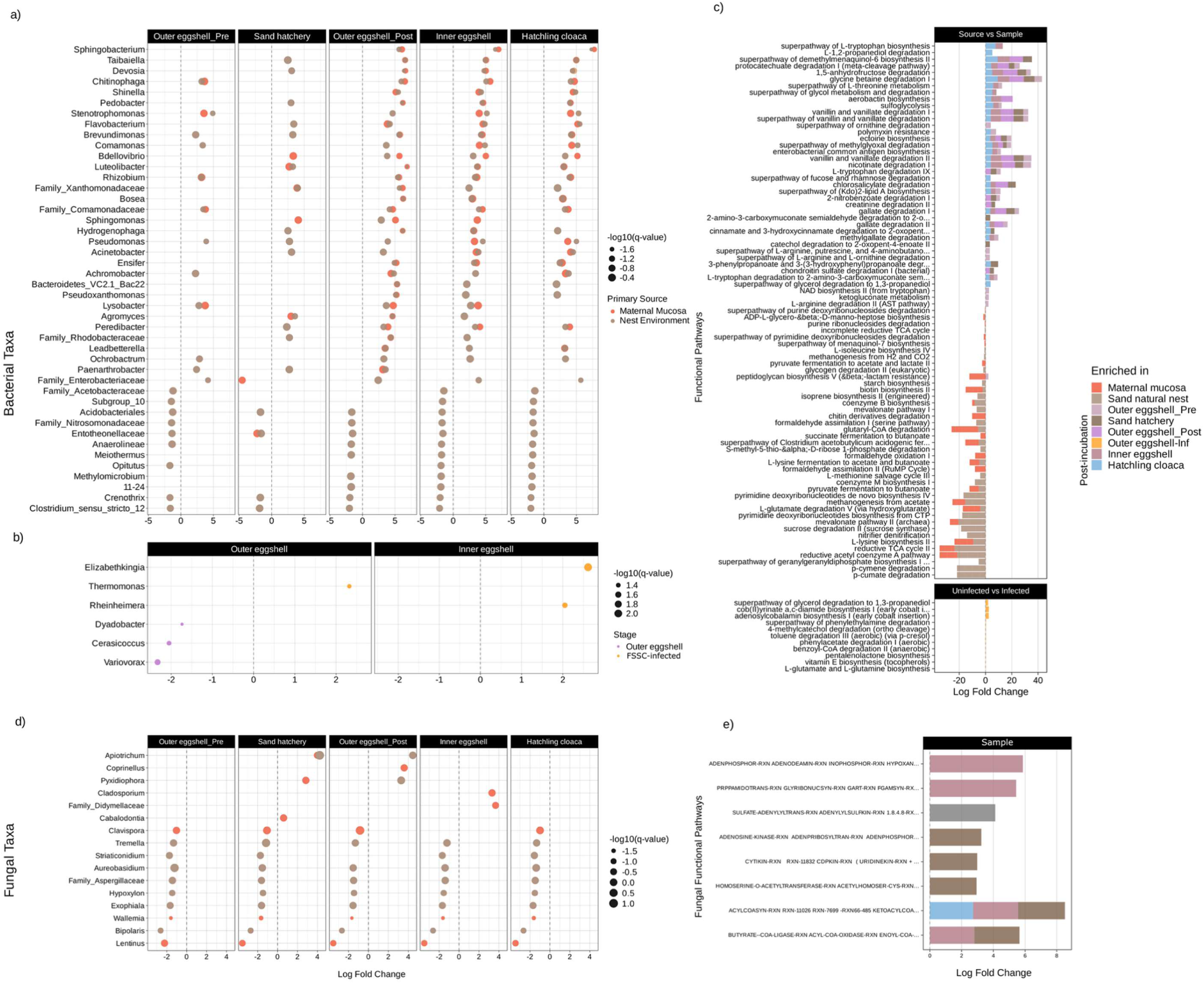
Differential abundance analysis of bacterial and fungal communities and their predicted metabolic functions across all sample types and infection status. (a) Dotplot showing differentially abundant bacterial genera (ANCOM, adjusted p < 0.005) in pre- (outer eggshell) and post-incubation (sand hatchery, outer-eggshell, inner-eggshell and hatchling cloaca) from bacteria consistently enriched in at least three sample types. To focus on biologically relevant changes, only genera with log₂ fold change ≥ -2 are displayed. Dot size indicates statistical significance (-log10(q-value)), and colour indicates the bacterial source (peach: maternal mucosa; light brown: nest environment). The x-axis shows log2 fold change, with negative values indicating enrichment in both maternal and environmental sources and positive values indicating enrichment in the different samples. (b) Barplots of differentially abundant MetaCyc pathways predicted from the bacteriome of pre- and post-incubation sample types. Bar length represents log2 fold change, with colours corresponding to bacterial source and the different sample types. For space reasons, 20 pathways that are significant in both sources and across at least three sample types are shown (ANCOM, adjusted p < 0.005). (c) Dot plot comparing differentially abundant bacterial genera between uninfected and fusariosis-infected samples in post-incubation samples (outer and inner eggshells). Positive log2 fold change indicates enrichment in fusariosis-infected samples. Dot size represents statistical significance, and colours distinguish uninfected and fusariosis-infected sample types. (d) Barplots of bacterial MetaCyc pathways enriched in fusariosis-infected samples (ANCOM, adjusted p < 0.005). (e) Dot plot showing differentially abundant fungal genera (ANCOM, adjusted p < 0.05) across pre- and post-incubation sample types, following the same visualisation approach as panel (a). (f) Bar plots of MetaCyc pathways predicted from the mycobiome, displaying functional differences across different sample types (ANCOM, adjusted p < 0.05).

When testing for infection, we did not identify significantly enriched fungal genera. However, we found that bacteria from the genus *Dyadobacter*, *Cerasicoccus* and *Variovax* were enriched in the outer eggshell of uninfected eggs (Fig. 5b), whilst bacteria from the genus *Thermomonas* were enriched in the outer eggshell of FSSC-infected eggs (Fig. 5b). Additionally, in the inner eggshell, bacteria from the genus *Elizabethkingia* and *Rheinheimera* were enriched in FSSC-infected eggs (Fig. 5b). No bacteria were enriched in uninfected inner eggshells.

We conducted Pycrust analysis to understand the functional profiles of bacterial and fungal communities. As with the taxonomic data, we used AMCOM-BC2 to test for differential abundance of MetaCyc functional pathways between pre- and post-incubation samples and between infection status groups. From these analyses, we observed several Metacyc pathways derived from bacteria in the pre- and post-incubation samples (Fig. 5c,e; Table S10a). Among these were pathways involved in amino acid biosynthesis (such as the superpathways of L-tryptophan biosynthesis, L-tyrosine biosynthesis, L-aspartate, and L-asparagine biosynthesis, among others; Table S10a), energy metabolism (including several TCA cycle variants, the superpathway of glycolysis, and pyruvate hydrogenase; Table S10a), carbohydrate metabolism (e.g., L-arabinose degradation, glycogen degradation, and the urea cycle), and cofactor and vitamin biosynthesis. These findings suggest that the bacteriome associated with incubation plays a key role in embryonic development and hatching success, as some of the listed functional pathways, such as the reductive TCA cycle and the reductive acetyl-CoA pathway, have previously been linked to the bacteriome of hatched eggs from this turtle species (25). MetaCyc pathways detected in the mycobiome were associated with amino acid biosynthesis, nucleotide metabolism and biosynthesis, fatty acid and energy metabolism, and nutrient assimilation. This suggests that the chemical makeup, dominated by fungi, may be linked to fungal pathogens or fungal growth (Fig. 5e, Table S10b). When assessing the impact of infection, we observed significant MetaCyc pathways linked to bacteria but not to fungi. Eleven bacterial MetaCyc pathways were enriched in FSSC-infected outer eggshells (Fig. 5c, Table S10c). Among these, the functions of the enriched pathways were primarily related to aromatic degradation (e.g., 4-methylcatechol degradation, toluene degradation III), cofactor biosynthesis (e.g., vitamin E biosynthesis, adenosylcobalamin biosynthesis I), and anaerobic metabolism (e.g., benzoyl-CoA degradation II). These results suggest the presence of pathogenic bacteria, which may be correlated with fungal infection and niche competition.

### Co-occurrence interactions between the most prevalent bacterial and fungal genera in relation to incubation and FSSC-infection

Microbial interactions change and shape the community throughout development, turning them into valuable indicators of community organisation, functional specialisation, and resilience (26). Here, we aimed to analyse interkingdom interactions (bacteria-fungi) in relation to incubation and fusariosis infection. We constructed microbial association networks to identify co-occurrence interactions between the 20 most prevalent bacterial and fungal genera using the Spearman measure and netAnalyze from the NetComi package. We compared positive and negative associations between pre-incubation (outer-eggshells) and post-incubation (outer-eggshells, inner-eggshells and hatchlings’ cloaca) samples and the effect of FSSC-infection among post-incubation samples.

Network analyses revealed distinct structural inter-kingdom organisation across pre- and post-incubation samples, ranging from 56 to 177 significant associations (r ≥ 0.3) (Fig. 6). Pre-incubation outer eggshells exhibited a complex network with 128 significant associations (density = 0.164) organised into five distinct modules (modularity = 0.332). However, no hub taxa dominated the community. Instead, centrality was distributed across multiple bacterial genera, particularly *Chitinophaga*, *Rhizobium*, and the family Intrasporangiaceae, each with high connectivity (15, 15, and 13 connections, respectively). This distributed structure suggests that outer eggshells host an established consortium of co-adapted microorganisms from maternal and nest environments.

**Fig. 6.**
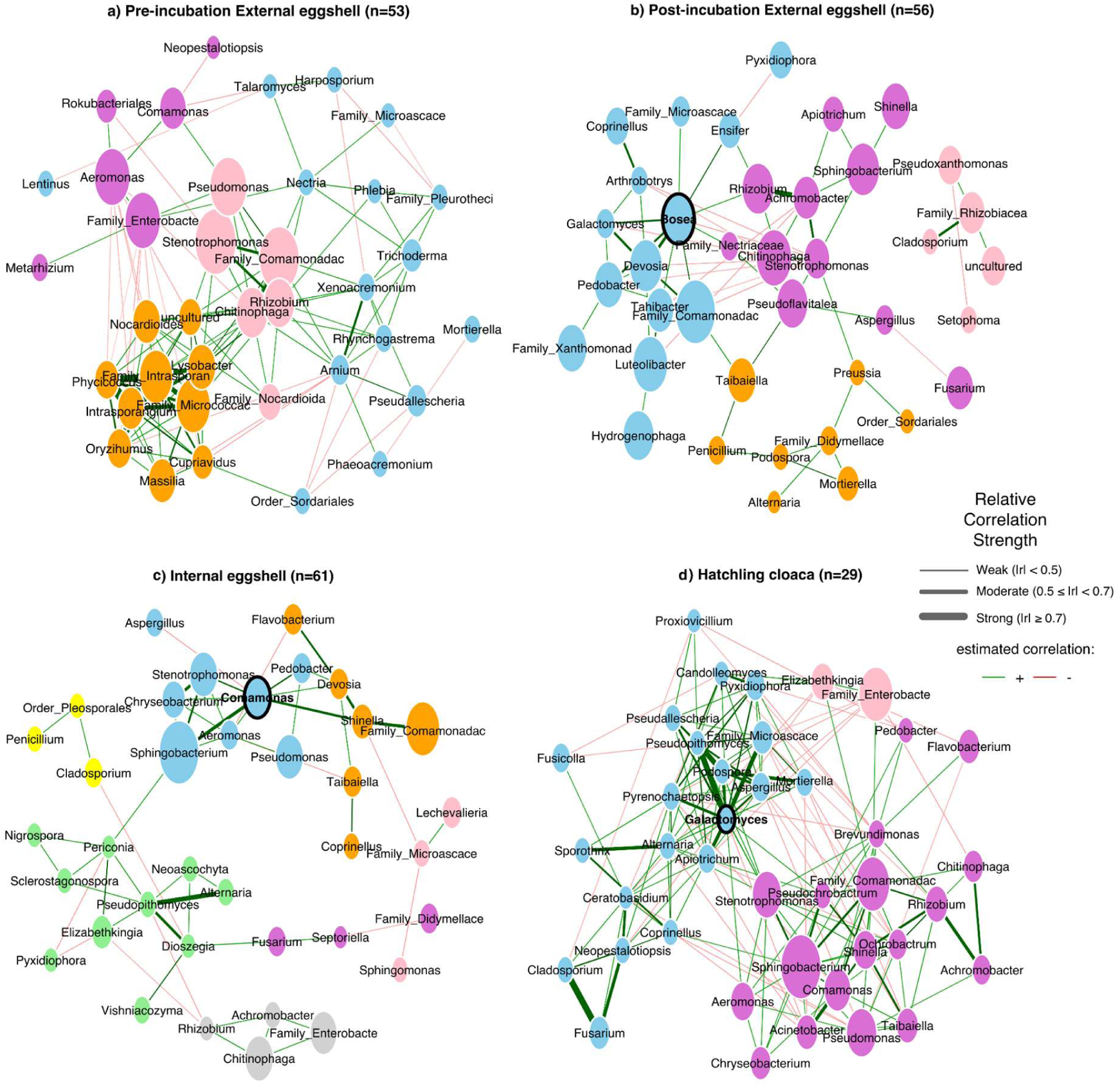
Co-occurrence networks showing interkingdom associations between the 20 most prevalent bacterial and fungal genera in pre- and post-incubation samples. a) Pre-incubation outer eggshell, b) post-incubation outer eggshells, c) inner eggshells, and d) hatchling cloaca. Networks were constructed based on Spearman correlations (r ≥ 0.3), with node size scaled to the sums of clr-transformed data and node colours representing the determined clusters. Single nodes were removed from the networks, with taxa labelled at the genus level. Edge thickness represents correlation strength. Hub taxa are highlighted in black, and n = X samples per group. A green line indicates positive correlations between fungi and bacteria, while a red line indicates negative correlations. Thicker lines represent stronger Spearman correlations.

After incubation, outer eggshells showed a more fragmented network (Fig. 6b) (modularity = 0.410), with only 65 associations (density = 0.083), a moderate correlation strength within the community (0.62), and a single hub, dominated by the genus *Bosea*. Inner-eggshells showed further network compartmentalisation with low network density (0.072) and only 56 associations distributed across eight modules (modularity = 0.576). The bacterial genus *Comamonas* emerged as the only hub, and the fungal genera *Alternatia* and *Pseudopithomyces* (r = 0.53) appeared among the top associations. In contrast, hatchling cloacal networks were more complex, with 177 associations and the highest edge density (0.227) among all sample types. This is reflected in the network’s connectivity, which is the only network to exhibit a fully connected community (modularity = 0.283), where every genus is directly or indirectly associated with all others. This suggests a tightly integrated community structure and functional integration, rather than competitive compartmentalisation. Notably, the hub taxa and the strongest associations shifted toward fungal-dominated genera, with *Galactomyces* emerging as the keystone genus, connecting to nearly half of the network nodes. The strongest associations include seven fungal pairs: *Cladosporium-Fusarium* (r = 0.76), *Galactomyces-Pseudopithomyces* (r = 0.74), and *Podospora-Pseudopithomyces* (r = 0.71). Bacterial genera that dominated pre-incubation eggshells (*Chitinophaga*, *Rhizobium*, and the family Intrasporangiaceae) were absent, replaced by *Stenotrophomonas* and *Sphingobacterium*. Overall, our results reveal a striking process in which microbial communities fragment during and after incubation, only to be replaced by a fundamentally new, integrated community in the hatchling’s cloaca. This suggests that microbial community dynamics are tissue-dependent and further supports the idea that host selection plays a critical role in community structuring.

Network communities showed different organisations in FSSC-infected samples. Outer eggshells exhibited distinct network structures depending on infection status (Fig. S3a). FSSC-infected outer eggshells showed more compartmentalised community (modularity = 0.268) with lower connectivity (edge density = 0.202) compared to uninfected eggs (modularity = 0.244; edge density = 0.264). However, positive associations increased from 67% to 76% in FSSC-infected external eggshells, suggesting a shift towards more cooperative interactions. Hub taxa shifted from two fungal genera (*Galactomyces* and *Alternaria*) in uninfected samples to the bacterial family Comamonadaceae in FSSC-infected outer-eggshells. Interestingly, in uninfected outer eggshells, *Fusarium* showed a strong negative association with *Aspergillus* (r = -0.76), suggesting competition between the pathogen and possible commensal fungi.

Inner eggshells showed a strong response to FSSC-infection (Fig. S3b), with FSSC-infected networks compartmentalised in three modules (modularity = 0.485) with lower edge density (0.144) compared to uninfected inner eggshells, which showed a more interconnected network (modularity = 0.277; edge density = 0.237). The genus *Comamonas* emerged as the hub taxa in the community. In contrast, centrality in uninfected samples was distributed across multiple taxa, suggesting that *Comamonas* may be involved in stress-tolerant interactions that bridge microbial groups isolated by the infection. Interestingly, *Fusarium* did not show antagonistic interactions in uninfected inner eggshells but formed the strongest association with the fungal genus *Neoascochyta* (r = 0.83), suggesting the formation of saprophitic consortia. In FSSC-infected inner eggshells, the strongest correlations were dominated by the fungal genera *Alternaria* and *Pseudopithomyces* (r = 0.68) and the bacterial genera *Comamonas* and *Shinella* (r = 0.62). These results suggest that infection promotes the growth of other pathogens and highlights resistance in the bacterial commensals.

In stark contrast to the eggshells, the uninfected hatchlings’ cloaca showed a less complex network compared to the FSSC-infected hatchlings’ cloaca (Fig. S3c). The network community of FSSC-infected hatchlings’ cloaca showed less compartmentalisation (modularity = 0.116) and increased connectivity among taxa (edge density = 0.467) compared to uninfected networks (modularity = 0.312; edge density = 0.271). However, positive associations were higher in uninfected hatchlings’ cloaca (82%) compared to FSSC-infected hatchlings’ cloaca (63%), suggesting increased competitiveness or antagonistic interactions in FSSC-infected individuals. Additionally, the hub in the uninfected hatchlings’ cloaca network was dominated by the bacterial genus *Shinella*. In contrast, there was no dominant hub in the cloacal network of FSSC-infected hatchlings; centrality was distributed across multiple highly connected taxa. Interestingly, *Fusarium* showed strong interactions in both uninfected and FSSC-infected cloaca networks with the bacterial genus *Cladosporium* (r = 0.73; r = 0.930, respectively), suggesting that *Fusarium* is not isolated from or excluded by the community, but rather becomes integrated into the developing cloacal microbiome. In uninfected networks, the second-strongest interaction was between the bacterial genera *Comamonas* and *Sphingobacterium* (r = 0.714), while in the FSSC-infected networks, the fungal genera *Galactomyces* and *Pyrenochaetopsis* exhibited the strongest interaction (r = 0.959). Our results suggest that microbial communities in different tissues respond differently to FSSC infection; specifically, those in the inner eggshells appear fragmented. In contrast, the microbial community in the hatchling’s cloaca remains robust despite FSSC infection. However, it is important to note that the sample size in the FSSC-infected hatchlings’ cloaca was smaller than in the uninfected cohort. Overall, our results indicate that FSSC infection alters microbial community structure, but host filtering plays a key role in selecting commensal microbes that help to stabilise the community.

## Discussion

Understanding microbial assembly during early life is crucial in microbial ecology. Particularly, the mechanisms by which hosts acquire their microorganisms - a process known as holobiont inheritance- remain an open question (26, 27). The nidobiome concept posits that environmental and maternal sources, along with host selection, play integral roles in shaping microbiome assembly throughout development (12). However, for oviparous vertebrates that lack parental care and develop in microbial-rich environments, our understanding of the mechanisms governing host-microbial assembly and their contributions to healthy development remains limited. In this study, we investigated whether the nidobiome concept applies to oviparous vertebrates without parental care and examined the role of host selection in microbial assembly. We also assessed the impact of fusariosis infection on microbiome assembly post-incubation. Our findings indicate that host filtering of maternal and environmental microbes helps beneficial symbionts to reach protective thresholds, enabling hosts under stress to retain a broad repertoire of microbes that defend against disease. This mechanism supports the resilience of turtle offspring microbiomes during fusariosis infection and promotes successful hatching.

### Vertically transmitted microbiome influences the nest microbiome, and subsequent host microbial assembly

We observed that at oviposition, maternal mucosa and outer eggshells exhibited similar bacterial species richness, whereas nest sand contained significantly higher and distinct bacterial diversity. This pattern suggests that maternal and environmental sources contribute different microbial inocula, with maternal microbes being successfully attached to the eggshell at oviposition. In vertebrates, including turtles, maternal mucosa produces glycosylated mucins that form a dynamic gel, acting simultaneously as a selective barrier, attachment surface, and nutrient source for microbes (28), likely playing a central mechanistic role in eggshell microbial inoculation. Supporting evidence from other oviparous reptiles, such as lizards and freshwater turtles, shows that eggshell communities more closely resemble maternal cloacal than oviductal microbial communities (16, 17).

Based on the hypothesis that early-arriving microbes (e.g., via vertical transmission) might influence the assembly of later-arriving microbes (e.g., via horizontal transmission) (13), we observed contrasting patterns in eggshell and nest bacterial communities after incubation. Outer eggshells retained their initial microbial richness but exhibited increased evenness, indicating community restructuring without the addition of new taxa. In contrast, nest sand showed an increase in bacterial richness without changes in evenness, suggesting a stable community structure with the accumulation of rare taxa. These observations suggest that vertical transmission establishes priority effects, where maternally provisioned microbes colonise eggshells before environmental microbes arrive, potentially modifying the environment in a way that influences the establishment of later-arriving microorganisms (i.e. niche facilitative modification) (13, 29). Comparable nest-host microbial dynamics have been observed in the Oriental Tit (*Parus minor*) during incubation (9) and in other passerine birds, where maternal cloacal communities influence both nest and offspring microbiomes (30). The persistance of bacterial richness on eggshells despite continuous environmental microbial influx indicate that vertically transmitted communities become established early and resist displacement by environmental colonisers, consistent with priority effects that shorten transient dynamics and reduce colonisation by later-arriving taxa, including potential pathogens (26, 31).

SourceTracker analyses further support our diversity results and reinforce the hypothesis that both vertically and horizontally transmitted microbial colonisers shape host community structure. Across sample types - from the maternal cloaca to the hatchling cloaca - forty-one bacterial and nine fungal taxa were shared, indicating that maternal and environmental sources both play significant roles in microbial community assembly. These findings suggest that microbial communities from the mother and the environment are transferred to eggshells and hatchlings’ cloaca, consistent with a mixed mode of microbial transmission, as observed in freshwater turtles (17). Given that priority effects favour early-arriving microbial colonisers (13, 31), and as described above, maternal microbes appear well-established on eggshells at oviposition, we propose that maternal inocula not only seed the eggshell microbiome but also shape the surrounding nest microbiome, creating conditions that facilitate host microbial assembly.

### Communities converge toward a consistent functional microbiome composition through active host selection

After incubation, we observed that bacterial and fungal diversity of inner eggshells and hatchlings’ cloaca were significantly lower than on outer eggshells, yet similar to each other, while microbial composition was distinct in each of the host compartments (i.e. eggshells and hatchlings’ cloaca). Moreover, community dispersion progressively decreased from heterogeneous outer eggshells to the more homogeneous inner eggshells and hatchling cloaca, indicating deterministic rather than stochastic assembly, consistent with host-genetics-driven microbiome assembly in fish embryos (32, 33).

Differential abundance analyses strongly supported the presence of priority effects and host-mediated selection processes in microbial assembly. Bacterial taxa enriched across outer eggshell, inner eggshell, and hatchling cloaca—including *Bdellovibrio, Brevundimonas, Devosia, Ensifer, Flavobacterium, Luteolibacter, Lysobacter, Pseudomonas, Peredibacter, Sphingomonas, Sphingobacterium, Taiabella, Shinella,* and *Acinetobacter*—have been previously reported as part of the commensal eggshell microbiota of this turtle species and other oviparous vertebrates (11, 25). Critically, all these genera are known to produce antifungal metabolites that suppress *Fusarium oxysporum* and other pathogens in the rhizosphere, and have been linked to suppression of fusariosis disease and healthy hatching outcomes in this species (25, 34–37). The ongoing presence of essential taxa in various host tissues, both before and after incubation, underscores the influence of host selection. *Acinetobacter, Sphingomonas, Sphingobacterium, Chitinophaga, Bosea, Lysobacter,* and the family Comamonadaceae were enriched in both eggshells and hatchling cloaca. *Sphingobacterium* has been previously detected in the cloacal microbiota of freshly hatched *P. unifilis* turtles and in the gut microbiome of amphibians (11, 38, 39), while *Lysobacter, Comamonas,* and *Bosea* persisted throughout hatchling development for over 25 days post-hatching (11). The presence of these taxa in pre-incubation outer eggshells confirms that maternal transmission, environmental influence, and host selection jointly shape microbial assembly from oviposition onward, establishing a functional microbiome that persists through early development. Research in amphibians and reptiles suggests that gut microbial communities may have co-evolved with their hosts, retaining core bacterial taxa that enhance development and protect against fungal infections (40, 41). However, studies on herpetofauna microbiomes remain limited.

PICRUSt2 functional predictions further supported the developmental significance of these microbial communities (Table S10). Superpathways of polyamine biosynthesis I and II, which modulate developmental arrest in *C. elegans* embryos with TCA cycle deficiency, are known to stimulate embryogenesis and ensure developmental progression (42). The TCA cycle itself, detected consistently across tissues, including the healthy inner eggshells of this species (25), ensures mitochondrial function during embryogenesis by regulating glycolysis and fatty acid synthesis (42) and controls transcriptional activation and epigenetic remodelling in mammalian embryos (43). Carbohydrate biosynthesis pathways, particularly those involved in amino acid metabolism and arginine biosynthesis, were also detected and align with their reported roles in promoting growth and mitigating oxidative stress during late embryonic development in birds (44). Furthermore, the presence of short-chain fatty acid pathways (e.g., succinate fermentation, reductive acetyl-CoA pathway) and vitamin K2 biosynthesis pathways (e.g., superpathway of demethylmenaquinol-8 biosynthesis) suggests microbial contributions to calcium metabolism during incubation (45, 46). This is particularly relevant in turtles, where calcium is progressively absorbed from softening eggshells and the yolk throughout development (47, 48). Collectively, these results provide compelling evidence that host selection plays a critical role in microbiome assembly by filtering maternally and environmentally transmitted microbiomes that have significant functional roles in development and health.

Further supporting our results, we observed that the microbial communities undergo substantial restructuring during incubation. Pre-incubation outer eggshells exhibited a distributed bacterial network with no dominant hub taxon, suggesting that the maternal cloacal mucosa harbours an established consortium of co-adapted microorganisms. While co-occurrence networks in microbial ecology are limited and conclusions must be cautious, dropout experiments with synthetic microbial communities suggest that microbial interactions in the absence of hub taxa can still maintain functionality (49). Strong correlation among bacterial genera such as *Chitinophaga, Rhizobium,* and *Intrasporangiaceae* likely represents an environmental signature successfully colonising egg surfaces during oviposition, as *Rhizobium* and, especially, *Chitinophaga* appear to originate from the environment. This supports the idea that vertically transmitted microbial communities are well established during oviposition. When vertical transmission predominates, the microbiome can reach steady-state density before horizontally acquired microbes have a substantial influence (29), suggesting that vertical transmission is a stronger force than horizontal acquisition in oviparous vertebrates lacking parental care. After incubation, outer eggshell communities became more compartmentalised, with the genus *Bosea* emerging as a hub taxon, while inner eggshells showed further compartmentalisation with *Comamonas* serving as the hub. This indicates that eggshells develop a complex and functionally stable microbial community during incubation, likely mediated by host selection and host genetics, as observed in the gut microbiomes of other systems, including humans, fish, and mice (3, 50). Moreover, physical confinement of microbes to specific host tissues can mediate microbial interactions and even drive the evolution of metabolic dependencies between taxa (51, 52). Together, these findings suggest that host filtering actively shapes microbiome composition throughout development.

In stark contrast to the fragmented eggshell networks, hatchling cloacal communities exhibited the highest connectivity of all sample types. This tightly integrated architecture indicates functional coordination rather than competitive compartmentalisation, a hallmark of mature, host-associated microbiomes (52, 53). Notably, the cloacal microbial structure differed from that of the eggshells: while bacteria predominated on eggshells, the cloacal environment was dominated by fungal hubs and strong inter-fungal connections. *Galactomyces* emerged as a key player, linking nearly half of the network nodes. The bacterial taxa present on eggshells before incubation, *Chitinophaga*, *Rhizobium*, and *Intrasporangiaceae*, were absent in the hatchling cloaca, which instead featured *Stenotrophomonas* and *Sphingobacterium*. This shift indicates that the host actively selects its microbiota rather than passively inheriting it from the eggshell. The combination of eggshell–derived bacteria and nest–sand–derived fungi produces a diverse and functionally effective community, as reflected by the high connectivity among various fungal genera. These results suggest that the host selectively recruits symbionts from the available pool to enhance host development and overall fitness. It is important to recognise that our findings are based on metabarcoding, which provides a snapshot of community composition but does not capture functional interactions or confirm whether host filtering actively modifies symbiont interactions. Addressing these questions will require complementary meta-omics approaches.

### FSSC infection alters microbial assembly process, yet the host filters for protective microbial taxa

Host control over microbiota is expected to enhance fitness benefits either by altering which symbionts are present (i.e. partner choice) or by changing the symbiont phenotype (i.e. partner manipulation) (3). Such control can manifest not only in community composition but also in the associations among coexisting microbial taxa during host development (54). In this study, fusariosis infection was first detected after incubation and was associated with significant shifts in both outer and inner eggshell bacterial communities, while hatchlings’ cloaca communities remained comparatively stable. Sourcetracker analysis further revealed that FSSC-infected individuals shared fewer microbial taxa (36 bacterial and five fungal genera) than uninfected individuals (41 bacterial and nine fungal genera), suggesting that FSSC pathogens may disrupt both bacteriome and mycobiome community assembly processes. Notably, *Fusarium* was detected in both uninfected and infected samples, and across both maternal and nest sources. This pattern indicates that *Fusarium* is likely widespread in the environment, consistent with previous studies (23, 55). Additionally, the presence of FSSC members in maternal cloaca supports the idea that the maternal cloaca may act as a reservoir of potential pathogens, as reported in the sea turtle *Caretta caretta* (23). Whether *Fusarium* pathogenicity is determined by host species identity or by specific host or environmental conditions remains an open question.

Interestingly, during microbial transmission, bacterial genera such as *Taiabella*, *Stenotrophomonas*, *Sphingobacterium*, *Pseudomonas, Flavobacterium, and Chitinophaga* were more abundant in uninfected individuals than in those infected with FSSC. As discussed above, these taxa form part of the commensal bacterial community of *P. unifilis* inner eggshells (11, 25); specifically, *Sphingobacterium* and *Pseudomonas* have been directly linked to hatching success and FSSC infection resistance in this species (25), while *Flavobacterium* has been reported as a commensal in the gut microbiome of the Chinese soft-shelled turtle (17). Moreover, bacterial genera such as *Stenotrophomonas*, *Pseudomonas*, and *Flavobacterium* possess well-characterised antifungal activity against *Fusarium spp.* and other fungal pathogens (56–58). Therefore, their collective decline during infection indicates a disruption of microbes that may be involved in antifungal defence. Interestingly, two bacterial genera, *Paenarthrobacter* and *Bosea*, were detected exclusively in infected individuals. Bacteria in the genus *Paenarthrobacter* are known to produce volatile compounds that induce oxidative stress and disrupt the integrity of fungal cell walls, including those of *Fusarium spp.* (39, 59). *Bosea* has been associated with antimicrobial resistance genes and antifungal activity specifically against *F. proliferatum* and *F. oxysporum* (60). Together, the presence of these bacterial taxa in infected individuals suggests that during assembly, the microbial community offers an antifungal response, yet this response appears insufficient to fully prevent pathogen establishment and community disruption. Indeed, enrichment of *Chryseobacterium* and *Morganella* in infected eggs indicates potential proliferation of pathobionts. While *Morganella* (family Enterobacteriaceae) occurs as a commensal in multiple tissues of *P. unifilis* and *P. expansa*, it can become pathogenic when the host is immunocompromised or when a primary infection is present (61), as may be the case here. In the fungal community, *Apiotrichum* was predominant in uninfected samples and derived primarily from the nest environment. This genus encompasses saprotrophic species with roles in decomposition and nutrient cycling, though recent evidence shows that some species can act as opportunistic human pathogens by growing at human body temperature, a physiological barrier that typically restricts fungal pathogenicity (62). Whether *Apiotrichum* represents a commensal component of the nest fungal community or poses a risk to embryo development in *P. unifilis* cannot be resolved from the present data. *Coprinellus* was detected in both infected and uninfected individuals but showed a higher prevalence in infected inner eggshells, consistent with previous observations in this species (25). As a saprotrophic fungus, *Coprinellus* may function as a commensal taxon involved in the decomposition of organic matter released during hatching. Its predominance at this stage, rather than in earlier developmental samples, further supports this interpretation.

While FSSC infection seems to alter outer and inner eggshell composition, hatchling cloaca communities remained unaffected. Moreover, *Fusarium* abundance in the hatchling cloaca did not differ markedly between infected and uninfected individuals. This may indicate that successfully hatched individuals overcome the infection, facilitated by antifungal bacterial and fungal communities from both maternal and environmental sources. This is consistent with experimental evidence in lizards showing that vertically transmitted bacteria can suppress fungal disease and improve hatching success (16), and with our previous findings that hatchlings that emerge despite infection harbour distinct, potentially protective bacterial communities (25).

Interestingly, FSSC infection revealed tissue-dependent network responses, highlighting contrasting vulnerabilities and resilience mechanisms. Inner eggshell networks underwent pronounced fragmentation, forming three disconnected subnetworks. Within these, *Comamonas -* a genus commonly found in soils and known to include species with antifungal activity against *Aspergillus* and *F. oxysporum* (63) - emerged as the sole hub taxon. This pattern suggests a strong environmental influence, whereby nest-sand-derived bacteria may act as connectors bridging communities otherwise isolated under pathogen pressure, consistent with studies showing that hub and connector taxa structure microbial network communication and stability in soil systems (64, 65). At the same time, *Fusarium* formed saprophytic fungal consortia, establishing pathogen-dominated modules rather than displacing resident taxa. In contrast, cloacal networks exhibited intensification rather than disruption, maintaining a highly interconnected network structure. Here, *Fusarium* became integrated into the network, forming strong positive associations with resident fungi, such as *Cladosporium*, indicative of pathogen integration rather than exclusion. The observed trend towards more competition-dominated resource partitioning (positive associations decreasing from 82% → 63%) may reflect maturation towards more stable and established microbiomes capable of absorbing perturbations. This suggests that host health may depend not only on community composition but also on the underlying network structure. These findings align with the hypothesis that host selection can influence symbiont behaviour and phenotype, potentially promoting cooperative or even altruistic traits within the microbiome (i.e. the holobiont) (3, 29). Moreover, ecological theory predicts that imperfect vertical transmission may be advantageous, allowing hosts to acquire beneficial symbiont combinations suited to current environmental conditions (66). Such patterns have been documented in diverse systems, including corals (67, 68) and fungus-farming termites (69). Another important protective mechanism is colonisation resistance, whereby resident microbiota occupy ecological niches that might otherwise be exploited by pathogens (70, 71). Disruption of this mechanism, for example, through antibiotic treatment, can increase susceptibility to infection, whereas restoration of commensal communities can re-establish protection (70). Functional redundancy within microbial communities is a hallmark of resilient ecosystems (20) and may help explain the observed stability of cloacal microbiota under pathogen pressure. However, these findings should be interpreted cautiously due to the limited number of infected hatchlings.

Overall, our findings demonstrate that maternal transmission mediates the initial seeding of the egg microbiome, while the nest environment provides additional microbial inputs, consistent with previous work in this species (11). We show that host selection across development—acting on both maternally and environmentally derived microbes—shapes microbial community composition and symbiont interactions. These results extend the nidobiome concept to oviparous vertebrates lacking parental care, suggesting that maternal provisioning establishes priority effects, nest and eggshell communities undergo facilitative interactions during incubation, and deterministic host filtering progressively assembles more homogeneous offspring microbiomes from heterogeneous nidobiome sources. Notably, host-mediated microbial assembly remains robust under pathogen pressure, indicating that host-driven filtering, rather than environmental stochasticity or pathogen presence, governs early microbiome establishment in this system. Our results provide the first evidence in an oviparous vertebrate lacking parental care that maternally derived microbiota from the cloacal mucosa colonise both the egg and the nest environment, where they are subsequently shaped by host and environmental factors throughout development. Given the increasing incidence of bacterial and fungal diseases affecting turtle eggs globally (22, 23, 72, 73), understanding microbiome assembly and host selection may inform microbiome-based conservation strategies.

## Materials and Methods

### Study species and study site

The yellow-spotted Amazon freshwater turtle (*Podocnemis unifilis*) is found throughout the northern Amazon region, in the Orinoco, Amazon and Essequibo river basins, and the eastern Guianas. This species is threatened by over-harvesting across much of its geographic range. Details on the breeding ecology of *P. unifilis* are given in ref (74). Since 2007, a reintroduction project for *P. unifilis* has been established at the Tiputini Biodiversity Station (TBS), managed by the Universidad San Francisco de Quito, in the Yasuni Biosphere Reserve, Orellana Province, Ecuador (0°38′18″S 76°9′0″W). To stabilise and maintain the population, approximately 700 eggs are collected each year from nests along the banks of the Tiputini River and hatched in artificial nests at the research station. The turtles are kept in artificial ponds filled with river water for one to two months before being released back into the Tiputini River.

### Sample collection

For our project, we partnered with TBS to conduct a longitudinal study on the vertical (maternal) and horizontal (environmental) transmission of microorganisms to the egg and hatchling cloaca of *P. unifilis*. We aimed to explore the role of the nidobiome in influencing host fitness, particularly hatching success, in the context of stressors such as emerging *Fusarium* fungal pathogens. In 2022 and 2023, we collected egg and environmental samples during two key periods: before incubation (pre-incubation, laying season, December 2022) and after incubation (post-incubation, hatching season, March-April 2023), during which we also collected hatchling cloacal swabs.

During the pre-incubation phase, a total of 18 clutches were collected from natural nests, estimated to have been laid less than one day prior. After removing the sand covering the clutches, we collected mucosa from the female turtle’s cloaca that covered the eggs (hereafter referred to as maternal mucosa) using a sterile swab, taking care to avoid contact with the eggs’ surface. We then prepared a plastic box to facilitate the transfer of the eggs into artificial hatcheries. This box was filled with sand from the beach where the nest was located, and the eggs were covered with the same sand that had been in contact with the clutches, preserving the natural sand to the greatest extent possible. During clutch collection, we randomly selected 10 eggs from each clutch, rinsed them with distilled water, and swabbed the outer eggshell for up to 15 seconds (hereafter referred to as the outer eggshell, pre-incubation). Each swabbed egg was assigned a number from one to ten using a marker. Additionally, three sand samples were collected from each natural nest (hereafter referred to as sand natural nest): one from the sand covering the nest (the sand atop the clutch), one from the sand in direct contact with the clutch (surrounding sand once the eggs became visible), and one from the sand at the bottom of the clutch (taken from the sand surrounding the base of the nest after all eggs were removed). At TBS, the clutches were carefully transferred into wooden hatcheries, and two additional sand samples were collected from the artificial nest (designated as sand hatchery, pre-incubation): one from the bottom of the nest, composed of beach sand, and one from the sand surrounding the clutch, a mixture of the natural nest sand and beach sand.

After the 90-day incubation period (the expected incubation time for *P. unifilis* under artificial conditions), when eggs started to hatch, eggs were examined for FSSC infections, and samples were collected for later analysis (post-incubation phase). Initial assessment of FSSC fungal infections was conducted via visual inspection: hatched eggs were classified as symptomatic for fusariosis if they exhibited atypical coloured spots (e.g., green, pink, or greyish), while those without such markings were classified as asymptomatic. All egg samples were later tested by PCR for fusariosis using *Fusarium*-specific primers and target sequencing, as described in ref (22, 25).

The samples collected for the post-incubation phase included both the external (hereafter outer eggshell, post-incubation) and internal surfaces of successfully hatched eggs (hereafter inner eggshell, post-incubation). Prior to sampling, the eggshells were rinsed with distilled water to remove any sand particles from both surfaces, and the surfaces were swabbed repeatedly for up to 15 seconds immediately after the hatchlings emerged from the eggs. The inner eggshell samples contained residual albumen. After collecting the eggshell samples, the hatchlings were rinsed with distilled water, and cloaca swabs were obtained. Each hatchling was then marked, weighed, and the measurements (length and width) of both the carapace and plastron were recorded. Additionally, two samples were collected from the artificial nests (hereafter referred to as sand hatchery, post-incubation). One sample was taken from the upper section of the nest, and the other one from the lower section after the entire clutch had hatched.

After collection, each swab sample was placed in 1.5 µl Eppendorf tubes containing Nucleic Acid Preservation (NAP) buffer. This buffer is a mixture of salts, including EDTA disodium salt dihydrate, sodium citrate trisodium salt dihydrate and ammonium sulfate mixed with ultra-purified molecular water (75). The NAP buffer was prepared in the molecular laboratories of the Institute of Evolutionary Ecology and Conservation Genomics at Ulm University, Germany, following the instructions outlined in ref (76). It has been shown that the NAP buffer preserves DNA material for at least ten months at room temperature (76) and successfully preserves nucleic acids in microbiome studies (75, 77, 78). Following fieldwork, all samples were transported to the Institute of Evolutionary Ecology and Conservation Genomics at Ulm University, Germany, and stored at 4 °C until sequencing.

### Assessment of inner egg bacteriome, mycobiome and FSSC pathogens using Illumina high-throughput sequencing

We extracted DNA from swab samples following standard protocols for the NucleoSpin® 96 Soil extraction kit (Macherey-Nagel, Germany). Samples were eluted with 70 μl preheated SE buffer. For all the samples, we performed a two-step PCR. In the first-step PCRs, we targeted the V4 region of bacterial 16S ribosomal RNA (16S rRNA) to investigate the bacteriome using primers 515F (5’-GTGCCAGCMGCCGCGGTAA-3’) and 806R (5’-GGACTACHVGGGTWTCTAAT-3’) (79). To characterise the fungal mycobiome, we amplified the internal transcribed spacer (ITS) region using primers ITS1F (5’-CTTGGTCATTTAGAGGAAGTAA-3’) and ITS4R (5’-TCCTCCGCTTATTGATATGC-3’) (80, 81). Additionally, we tested for the presence of *Fusarium* species by amplifying a 430 bp region of the translation elongation factor gene (TEF1-alpha) using primers Fa_150 (5’-CCGGTCACTTGATCTACCAG-3’) and Ra-2 (5’-ATGACGGTGACATAGTAGCG-3’) (82). All primer pairs were extended with universal sequences CS1 (ACACTGACGACATGGTTCTACA) + 4N and CS2 (TACGGTAGCAGAG-ACTTGGTCT) for forward and reverse primers, respectively (Standard Biotools, USA), which served as adapters in the second PCR step.

In the first step of PCR, we had a total volume of 10 µl, composed of 1 µl extracted DNA, 1.5 µl pooled forward and reverse primers (0.3 µM), 5 µl AmpliTaq Gold™ 360 Mixmastermix (Applied Biosystems, USA) and 2.5 µl ultrapure dH_2_O. The first step PCR for the 16S rRNA V4 region was run under the following conditions: initial denaturation at 95 °C for 10 min; 30 cycles including denaturation at 95 °C for 30 s, annealing at 60 °C for 30 s, and elongation at 72 °C for 45 s; final elongation at 72 °C for 10 min. The first-step PCR for the ITS region was performed with an initial denaturation at 95 °C for 10 min; 30 cycles, including denaturation at 95 °C for 30 s, annealing at 54 °C for 30 s, and elongation at 72 °C for 60 s; and a final elongation at 72 °C for 10 min. The first-step PCR for the TEF-alpha gene was conducted with an initial denaturation at 95 °C for 10 min; 35 cycles, including denaturation at 95 °C for 30 s, annealing at 52 °C for 30 s, and elongation at 72 °C for 60 s; and a final elongation at 72 °C for 10 min. Finally, gel electrophoresis was performed on each sample to confirm PCR success.

The second-step PCR was carried out in 20 μl reactions in 96-well plates to incorporate Illumina sequencing adapters and a sample-specific barcode. A master mix containing 10 µl AmpliTaq Gold™ 360 mastermix (Applied Biosystems, USA) and 3.5 µl ultrapure dH_2_O was prepared. Each well was loaded with 13.5 µl mastermix and 4 µl of a unique barcode (Acces Array™ Barcode Library for Illumina Sequencers-384, Single Direction, Standard Biotools, USA), together with 2.5 µl of the first step PCR amplification used as an input template; this process was done for each of the markers used in this study. The barcoded samples were cleaned using the NucleoMag NGS Clean-Up and Size Select kit (Macherey-Nagel, Germany) with a 1:1 ratio of amplicons to beads on a Gene Theatre (Analytik Jena, Germany) following the manufacturer’s instructions. Clean amplicons were eluted to a final volume of 20 µl. The purity of the amplicons was controlled in a QIAxcel Advanced System (QIAGEN, Germany). The clean amplicons were then quantified with the Quantifluor® dsDNA System (Promega Corporation, USA, Madison) on a TECAN Infinite F200 Pro (Tecan Trading AG, Switzerland) following the manufacturer’s instructions. We subsequently pooled all samples to a final DNA concentration of 12 ng and diluted the pool to 6 nM. Finally, the library was loaded at 12 pM and sequenced on an Illumina MiSeq instrument using a 250 bp paired-end strategy with 10% PhiX being added to account for low diversity. In our sequencing library, we included three field sampling negative controls (tubes with NAP buffer brought to the field and opened without samples), six negative controls from DNA extraction controls and nine negative controls from the PCR. Hereafter, negative controls are referred to as “blanks”.

### Bioinformatic processing of the bacterial microbiome

All pre-processing steps were performed on a larger dataset encompassing multiple samples collected during the field seasons of 2022-2023 in a pristine environment and 2023-2024 in a disturbed environment. The analyses presented in this study are based on a subset of a dataset from the pristine environment, comprising all paired-end sequencing reads from 376 successfully sequenced eggshells, 67 hatchling cloacal, 11 maternal mucosa and 42 sand natural nest (from different parts of the 11 nests) and 43 (from different parts of the 11 nests) sand hatchery swab samples for the 16S V4 region were pre-processed by using the open-source QIIME2 microbiome analysis pipeline (version 2024.5) (83) and DADA2 (84) to remove dataset noise arising from artefacts and generate amplicon sequence variants (ASVs). Since error profiles differ across sequencing runs, DADA2 denoising was performed independently for each Illumina run to enable run-specific error-rate learning. The resulting ASV tables and representative sequences were subsequently merged across all runs prior to downstream analyses. We assigned taxonomy to the resulting 76,089 ASVs using the Silva (version 138.2) V4 (85) classifier as a reference. We removed sequences classified as chloroplast, mitochondria, archaea, Eukaryote, and unclassified phylum, resulting in 72,314 ASVs in the final dataset. Using MAFFT (86), we added an archaeal sequence to root a tree and constructed a phylogenetic tree using FastTree 2.1.8 (87). We exported QIIME2 tables to R and subsetted our data with the samples from the pristine environment indicated above for further analyses. After denoising, assigning taxonomic and filtering contaminants, we obtained 14’461,003 reads with an average of 24,427 per sample for downstream analyses. Contaminating bacterial DNA is commonly found in different buffers and extraction kits (88), and contaminants appear at higher frequencies in PCR controls than in positive samples (89). Therefore, we aimed to remove ASVs found in 51 successfully sequenced blanks (35 field blanks, seven extraction blanks and nine PCR blanks). We imported the data generated by QIIME2 using the R package “phyloseq” (90). In R, we used the function decontam::isContaminant (89) using the “prevalence” method to identify and remove the blank microbiome from the dataset. This filtering step resulted in the removal of 176 ASVs from the dataset. We excluded samples with fewer than 5,000 reads, removing six inner egg samples from our initial dataset. This left us with a final dataset that totals 14’284,272 reads across 30,109 ASVs and 588 samples, yielding an average of 24,292 reads per sample.

### Bioinformatic processing of the fungal mycobiome

All pre-processing steps were performed on a larger dataset encompassing multiple samples collected during the field seasons of 2022-2023 in a pristine environment. The analyses presented in this study are based on a subset of a dataset comprising all paired-end sequencing reads from 217 egg, 37 hatchling cloaca, 11 maternal mucosa and five sand natural nest and 19 sand hatchery (from different parts of 11 nests) for the ITS region, which were used to characterise the mycobiome. All paired-end sequencing reads were pre-processed using QIIME2 microbiome analysis pipeline (version 2024.5) (83) and DADA2 (84) in R using the DADA2 package to remove dataset noise arising from artefacts and generate amplicon sequence variants (ASVs). As the length variation of the ITS region (700 - 900 bp for this study) significantly impacts the filtering and trimming steps in the standard DADA2 workflow within QIIME2, we pre-processed the sequences in three different steps. First, we used the cutadatpt (91) plugin of QIIME2 to remove primers. Next, we extracted the ITS1 forward reads and ITS2 reverse reads (which encompass the entire ITS region) from the qzv. and save them into fastq. files. These files were placed in a new folder for further processing. Second, the ITS1 forward and ITS2 reverse reads without the primers were processed and concatenated with DADA2 through R and a fna. file was created to export the information to QIIME2 for taxonomic assignment. Third, we used the UNITE database release 10.0 2025 (92) for taxonomic assignment. Before building the classifier, we de-replicated the database using the dereplicate plugging of Rescript (93) in QIMME2. Dereplication is recommended to remove redundant sequence data from databases, as maintaining redundancy can cause sequencing reads to be randomly distributed across duplicate genomes. This leads to either a single random alignment reported from many possible options or multiple alignments at redundant locations. Such redundancy can skew the analysis of relative abundance, particularly for fungal diversity across samples, making it seem as if multiple ecologically equivalent populations coexist and resulting in an artificially low estimate of each taxon’s relative abundance (94).

After de-replication, a classifier was built, and taxonomy was assigned to 24,251 ASVs in the whole dataset. Using MAFFT (86), we added an opisthokonta sequence (*Fonticula alba*) to root a tree and constructed a phylogenetic tree using FastTree 2.1.8 (87). After denoising, assigning taxonomic, filtering contaminants and subsetting the data for the specific samples used in this study, we obtained 3’156,110 reads across 11,024 ASVs in 333 samples, with an average of 9,477 reads per sample for downstream analyses. Following the same approach as for the bacterial microbiome, we aimed to remove ASVs found in 31 successfully sequenced blanks (18 field blanks, 10 extraction blanks and three PCR blanks) using *decontam::isContaminant* (89) and the “prevalence” method. This filtering step resulted in the removal of 82 ASVs in the dataset. Finally, we excluded samples with less than 100 reads, resulting in a final data set with a total number of reads of 3’100,379 across 10,942 ASVs and 328 samples, with an average number of reads per sample of 9,969.

### Taxonomic assignment of *Fusarium* (FSSC) infections

All paired-end sequencing reads from 127 successfully sequenced samples (n= 106 eggs, n = 14 hatchlings’ cloaca and n= 7 nest sand) were pre-processed as described for the 16S and ITS markers. To assign taxonomy to 241 ASVs from the TEF1-alpha gene, we constructed a database from the NCBI Genbank using RESCRIPt (93) via QIIME2. We used the query “*elongation factor 1-alpha[All Fields] AND fungi[filter]*” to filter the database sequences by a minimum length of 100 and built a classifier. Before using the classifier for taxonomy assignment, we trained the databases with the specific TEF primers used in this study. Only sequences assigned to *Fusarium* or the teleomorph name (*Gibberella* and *Nectria*) longer than 300 bp were selected for further analysis. After denoising, assigning taxonomic and filtering contaminants, we obtained 97,640 reads across 28 ASVs and 125 samples, with an average of 769 reads per sample for downstream analyses. We included only samples with an FSSC ASV count greater than one, and incorporated this information as a variable in the overall bacteriome and mycobiome metadata.

### Statistical analysis

#### Variation of the bacteriome and mycobiome diversity across sample types collected before and after incubation, and the effect of infection

We investigated how bacterial and fungal diversity, as well as community structure, varied between pre-incubation (maternal mucosa, sand natural nest, outer eggshell) and post-incubation (sand hatchery, outer eggshell, inner eggshell, hatchling cloaca) sample types, and examined the effects of fusariosis infection status (FSSC-uninfected and FSSC-infected) on post-incubation biological samples.

We assessed alpha diversity using Observed species richness and the Shannon index. For observed richness, we applied generalised linear models (GLMs) with a negative binomial distribution, and for Shannon diversity we used a gamma distribution, both implemented in the R package “mgcv” (95). We compared two models: a null model including only FSSC infection status, and a full model including pre- and post-incubation samples and FSSC infection status as additive effects (sample type + FSSC). We assessed the significance of the sample type effect using likelihood ratio tests (LRT) for negative binomial models and F-tests for models with a gamma distribution. Post-hoc pairwise comparisons were performed using Tukey’s honestly significant difference (HSD) test using the “emmeans” package (96). To specifically test the effect of FSSC infection on post-incubation samples, we conducted separate analyses for these samples, including an interaction term (sample type × FSSC) to determine whether the infection effects varied across different post-incubation sample types.

We tested whether bacteriome and mycobiome composition are independently influenced by the incubation period (pre- and post-incubation) and FSSC infection status, or by the interaction of the two variables. To account for the compositional nature of microbiome data (97), we transformed abundance data using centred log-ratios (rCLR) as implemented in the “compositions” package (98). We performed redundancy analyses (RDA) using the “vegan” (99) with permutation tests (9,999 permutations) in separate models: 1) sample type effect across all pre- and post-incubation samples; 2) FSSC infection status effect in post-incubation samples only, and 3) interaction between sample type and FSSC infection status (sample type × FSSC) in post-incubation samples only. We visualised compositional differences using RDA ordination plots with Euclidean distances. Post-hoc pairwise comparisons between sample types were conducted using constrained RDA with Euclidean distances using the *“multiconstrained”* function from the “BiodiversityR” package (100).

We investigated whether the variation in bacteriome and mycobiome composition (heterogeneity) is associated with incubation or FSSC infection status. We predicted that heterogeneity of beta diversity (dispersion from the population median) would decrease from environmental samples to inner eggshells and hatchlings’ cloaca if deterministic host filtering processes shape microbial community assembly. We calculated beta dispersion using the *“betadisper”* function from the R “vegan” package (99) with Euclidean distances and median distance to the centroid within each sample type and infection status. To account for heterogeneity of variance between groups, we performed generalised least squares (GLS) models using the R package “nlme” (101). We compared models including sample type only, FSSC infection status only, and both variables (Sample type + FSSC) using AICc for model selection. We performed Tukey’s HSD tests from the best-fitting GLS model to test pairwise differences in dispersion between sample types and infection statuses.

#### Contributions of maternal and environmental microbes to the nesting environment and offspring

We conducted Sourcetracker2 (102) analyses to quantify the maternal and environmental contributions of microbes across all samples collected pre- and post-incubation and to test the effect of FSSC infection on microbial transmission patterns. The analyses included two sources (maternal mucosa and sand natural nest) and five sinks (pre-incubation: outer eggshells, post-incubation: sand hatchery, outer eggshell, inner eggshell and hatchling cloaca); uninfected and infected samples were analysed separately to avoid bias in the results. After importing the SourceTracker2 results into R, we selected taxa with a prevalence >10 in the bacteriome and >1 in the mycobiome, and taxa shared across all sample types. We then conducted intersection analyses to identify microbial sharing patterns across sample types, using UpSet plots generated with the R package “ComplexUpset” (103). Presence-absence matrices were constructed for microbial genera across all sample types, with genera filtered to include only those with non-zero abundance values. Sample types were ordered chronologically to illustrate temporal progression from pre-incubation to post-incubation stages. Additionally, we generated Sunburst plots using the R package “igraph” (104) and “plotly” (105) to visualise the taxonomic structure of microbial genera shared across all sample types. Sunbursts show taxonomic relationships from the phylum (inner ring) through family to genus (outer rings), with segment size representing each taxon’s relative abundance. To interpret microbial distribution and transmission across sample types, we used bubble plots, with bubble size representing relative abundance and bubble colour indicating the primary source as determined by SourceTracker analysis. Only genera present at key intersections in Upset analyses were included, representing the most prevalent and shared genera across development.

#### Changes in relative abundance of bacterial and fungal taxa and their associated MetaCyc metabolic pathways, according to incubation status and FSSC-infection

We estimated changes in the relative abundance of specific ASVs according to incubation period (pre- and post-incubation samples) and FSSC infection status in post-incubation samples, using Analysis of Compositions of Microbiomes with Bias Correction (ANCOM-BC2) (106). ANCOM-BC2 accounts for the compositional nature of microbiome data and corrects for sampling fraction bias, providing unbiased estimates of absolute abundance changes. In the incubation analysis, we compared microbial communities at five key time points (pre-incubation: outer eggshells, post-incubation: sand hatchery, outer eggshells, inner eggshells, and hatchling cloaca) against two reference sources: the maternal mucosa and natural nest environment. For the FSSC infection analysis, we compared uninfected versus FSSC-infected post-incubation samples. Taxa were included in the analysis if they were present in at least 10% of samples with a minimum abundance threshold of 10 reads.

Statistical significance was determined using the Holm-Bonferroni method for multiple testing correction, with an adjusted p-value threshold of 0.05. Log₂ fold changes were calculated to quantify the magnitude and direction of abundance differences: positive values indicate enrichment in the comparison group (pre- and post-incubation or infected samples), and negative values indicate enrichment in the reference group (maternal mucosa, sand natural nest, or uninfected samples). For visualisation of microbial genera abundance patterns, we applied a log₂ fold change threshold of ≥ -2 to focus on biologically relevant changes while excluding taxa with minimal depletion (< 4-fold decrease). Our analysis prioritised taxa that were significantly differentially abundant in both reference sources (maternal mucosa and nest environment) and present across at least three sample types, identifying core taxa consistently associated with the incubation process.

Functional metagenomic profiles were predicted from 16S rRNA gene sequences using PICRUSt2 (version 2.x) (107) and from ITS sequences using the PICRUSt2 pipeline through FROGSFUNC (108). MetaCyc pathway abundances were inferred and analysed for differential abundance using ANCOM-BC2, with the same statistical parameters applied as for taxonomic analysis. We selected pathways with significant differential abundance (adjusted p < 0.05) in both bacterial sources and across at least three sample types to identify core functional shifts associated with egg incubation and pathogen exposure.

#### Interkingdom co-occurrence networks before and after incubation and FSSC-infection

We performed Network analyses using the R package “NetCoMi” (109) to identify clusters of co-occurring interkingdom genera according to pre- and post-incubation sample types and FSSC infection status. As for beta diversity, we transformed abundance data using centred log-ratios (rCLR) to account for the compositional nature of microbiome sequencing data. The data were then filtered to retain the top 20 most prevalent bacterial and fungal genera using *microViz::tax_top()* (by = ‘prev’, n = 20). Pairwise associations were quantified using Spearman’s rank correlation, and only edges with a correlation coefficient |r|> 0.3 and a false discovery rate (FDR)-corrected P < 0.05 were retained to construct networks for each sample type and infection group. Separate networks were done for pre- and post-incubation outer eggshells, inner eggshells, and hatchling cloaca, as well as for uninfected and FSSC-infected subsets, to allow group-wise comparison of topological properties. Co-occurring clusters were identified using the *NetCoMi::netAnalyze* function with the fast-and-greedy method, and hub pars were calculated based on degree and eigenvector centralities. Clusters were visualised in different colours, and single clusters or taxa were excluded from the network.

## Acknowledgements

The authors are deeply grateful to all the staff of Tiputini Biodiversity Station, especially to Juan Carlos Rodriguez, Santiago Shiguango, José Macanilla, Mariano Grefa, and Luis Ganchoso, who assisted with field logistics, nest-finding and collection. Special thanks to Dr Alice Risely and Johannes Langpap for their assistance during field work. We thank the staff of the Plant Biotechnology lab at USFQ for all the support during fieldwork, Ciara Wirth for her assistance with the paperwork for the collection permits, Consuelo de Romo for her assistance with field logistics, Ulrike Stelle for her help with laboratory work, Dr Kunal Jani for sharing his opinion on the data analysis and Dr Mark Gillingham for his feedback and opinions on the study design and statistical analyses. The authors acknowledge the Kichwa Indigenous communities for granting access to their land and their efforts to conserve the Amazon forest and its biodiversity. The Tiputini Biodiversity Station and the Institute of Evolutionary Ecology and Conservation Genomics at Ulm University supported fieldwork This study was carried out under the collection research permit no. MAATE-ARSFC-2022-2399 and MAATE-DBI-CM--2023-0273 and exportation permits no.037-2024-EXP-CM-DBI/MAATE and no. 24EC000045E awarded by the Environment, Water and Ecological Transition Ministry of Ecuador. This study was funded by the German Research Foundation to S.S. (DFG SO 428/19-1), and A.S.C. was supported by a grant from the Gender Equality office during the preparation of the DFG proposal and fieldwork.

## Author contributions

A.S.C. conceived the idea, the study design, wrote the funding proposals, contributed to the issuing of fieldwork permits, contributed to fieldwork logistics, collected the data, completed all the laboratory analyses, prepared the samples for sequencing, analysed the data and wrote the manuscript; S.S. contributed to the study design, supervised the writing of funding proposal and was awarded funding, supervised the research project and gave feedback on the manuscript; K.W. contributed to the study design, supervised laboratory analyses and gave feedback on the manuscript. M.L.T. contributed to the study design, contributed to fieldwork logistics, supervised field work and gave feedback on the manuscript. D.R. contributed to the study design and fieldwork logistics and provided feedback on the manuscript. All authors gave final approval for publication of the manuscript, and there are no conflicts of interest.

